# Loofah, a newly characterized adhesion protein, suppresses cell death in long-lived *Drosophila* hindgut enterocytes

**DOI:** 10.1101/2025.04.04.647274

**Authors:** Jessica K. Sawyer, Ruth A. Montague, Olivia Goddard, Archan Chakraborty, Paulo B. Belato, Donald T. Fox

## Abstract

Tissue maintenance in the presence of cell death-promoting insults requires a host of molecular mechanisms. Many studies focus on cell renewal through regeneration, while fewer studies explore mechanisms that promote cell longevity despite cell death stimuli. Here, we reveal that the adult *Drosophila* hindgut ileum is an excellent model to study tissue maintenance by long-lived cells. Hindgut ileal enterocytes resist the damaging detergent SDS and upstream caspase signaling by *head-involution-defective (hid).* This *hid-*induced death insensitivity arises early in adulthood and associates with numerous transcriptional changes. We interrogated 82 of these transcriptional changes in a candidate screen for enhancers of *hid-*induced death in the ileum. Top among our screen hits is an immunoglobulin family cell adhesion gene, *CG15312. CG15312* maintains the adhesion protein FasIII on cell membranes. In *hid-*expressing ileal cells, *CG15312* loss causes cell death and pyknotic nuclear clustering. We name this conserved gene ***lo****w **o**n-membrane **f**<u>a</u>s and enhancer of <u>h</u>id* (*loofah*). Our findings reveal a new mechanism linking cell adhesion and cell death resistance in a long-lived cell type. Our work establishes a new model to study tissue preservation.

## Introduction

Tissue maintenance frequently requires regenerative mechanisms that replenish tissue mass that is lost over time (Poss and Tanaka 2024). Additionally, long-lived cells maintain tissues while lessening the burden on regenerative mechanisms (Wright et al. 2017; Guilliams et al. 2013; Zeps et al. 1996). Cellular and molecular mechanisms that distinguish long-lived cells from more transient cells remain poorly understood.

Intestinal tissue is exposed to the outer environment, and therefore tissue maintenance and regeneration or preservation strategies are critical. The *Drosophila* intestine is divided into three major organ segments: the foregut, midgut, and hindgut, each with distinct cell types and gene expression (Zhu, Ludington, and Spradling 2024; Marianes and Spradling 2013; Sawyer, Cohen, and Fox 2017; Cohen et al. 2018; Buchon et al. 2013; Miguel-Aliaga, Jasper, and Lemaitre 2018). Related to these differences in cell type, we previously noted a major tissue maintenance difference between the hindgut and midgut (Fox and Spradling 2009; Sawyer, Cohen, and Fox 2017). Whereas it had been established that the adult midgut epithelium contains actively dividing intestinal stem cells (ISCs) that replenish the midgut enterocyte (EC) population (Ohlstein and Spradling 2006; Micchelli and Perrimon 2006; Martin et al. 2018), the adult hindgut epithelium has minimal cell cycle activity and lacks active stem cells (Fox and Spradling 2009; Sawyer, Cohen, and Fox 2017). Therefore, regional differences in gene expression and stem cells coincide with distinct tissue maintenance biology of the *Drosophila* midgut and hindgut.

Even within the hindgut, tissue renewal mechanisms appear to differ. The hindgut epithelium consists of three regions: the pylorus, ileum, and rectum (Cohen et al. 2020). In this study, we focus on differences between the pylorus and ileum, and refer readers to our previous study of the rectum (Fox, Gall, and Spradling 2010; Schoenfelder et al. 2014; Stormo and Fox 2016, 2019; Peterson et al. 2020). The pyloric epithelium consists of small cells with little cytoplasm (Fox and Spradling 2009; Sawyer, Cohen, and Fox 2017). Unlike the pylorus, epithelial cells of the ileum exhibit properties of absorptive ECs, such as high expression of the Vacuolar H+ ATPase (Cohen et al. 2018), which is critical in transport across membranes, and Inebriated, a putative Na^+^/Cl^−^-dependent neurotransmitter/osmolyte transporter (Cohen et al. 2018; Luan, Quigley, and Li 2015). Pyloric cells possess regenerative properties. During metamorphosis, the larval ileum degenerates, whereas the pylorus generates new cells that comprise the adult pylorus and ileum (Cohen et al. 2021; Cohen et al. 2018; Fox and Spradling 2009; Takashima et al. 2008; Aghajanian et al. 2016; Zhang et al. 2024). In the adult hindgut, the regenerative potential of the pylorus is maintained, but the ability of the ileum to regenerate has been unclear. Our prior work employed an injury model whereby the pro-apoptotic genes *head involution defective* (*hid*) and/or *reaper* (*rpr*) are acutely induced by the Gal4/UAS system (see methods). Upon apoptotic cell loss of pyloric cells, the surviving pyloric cells enter the endocycle, thereby regenerating pyloric cell mass and genome content through increased ploidy and cell size (Cohen et al. 2018; Cohen et al. 2021; Losick, Fox, and Spradling 2013). Therefore, the adult pylorus retains the ability to regenerate the pyloric epithelium. However, in this model, apoptotic gene activation did not ablate ileal cells. Thus, whether the adult pylorus retains the ability to regenerate ECs of the ileum, much like midgut ISCs regenerate midgut ECs, remained an open question.

In this study, we reveal that hindgut ileal ECs are distinct from midgut ECs in multiple ways. Compared to midgut ECs, hindgut ileal ECs are long-lived and resist cell death from both ingested and genetically induced cell insults. In addition to possessing a protective apical cuticle, ileal ECs have distinct regulation of the apoptotic signaling pathway that promotes cell survival, namely resisting cell death by *hid* expression. However, these cells are highly sensitive to signaling downstream of *hid* by *death regulator Nedd2-like caspase (dronc)* caspase expression. This distinctive apoptotic regulation arises early in adulthood. By examining ileal gene expression differences during adult gut maturation, we conducted a candidate RNAi screen. This screen reveals an uncharacterized cell adhesion protein, that we name here **Lo**w **o**n- membrane **f**as and enhancer of **h**id (Loofah), is required for ileal cell insensitivity to *hid-* induced death. Loofah preserves the cell adhesion protein FasIII at ileal cell-cell junctions and prevents cell death upon *hid* expression. Loofah function appears critical in maintaining ECs of the hindgut, as unlike midgut EC cell loss, ablation of ileal ECs does not activate a robust regenerative response. Our findings underscore the importance of cell survival in a long-lived enterocyte population. Our results highlight a distinct long-lived, insult-resistant EC population in the *Drosophila* hindgut, and reveal a new tissue model and molecular mechanism to study the role of long-term cell survival in tissue maintenance.

## Results

### Adult hindgut ileum enterocytes are long-lived and resist the damaging agent SDS

Both the midgut and hindgut epithelium of adult *Drosophila* contain ECs, with all the hindgut ECs residing in the ileum (**Fig1A**, ileum ECs in green). Both types of ECs are distinguished by large polyploid nuclei relative to smaller non-EC nuclei. Adult midgut ECs are constantly renewed by intestinal stem cells (ISCs) (Ohlstein and Spradling 2006; Micchelli and Perrimon 2006), whereas adult hindgut ECs of the ileum arise during metamorphosis from an anterior region of the hindgut known as the pylorus (**Fig1B**)(Fox and Spradling 2009). Previously, we highlighted differences in cellular half-life between the adult ECs of the midgut and hindgut. Pulse labeling of actively replicating midgut and hindgut ECs with the thymidine analog Bromo deoxy Uridine (BrdU) suggested that hindgut ECs are much longer lived than midgut ECs (Fox and Spradling 2009). We later showed that, unlike the midgut (Ohlstein and Spradling 2006; Micchelli and Perrimon 2006), the adult hindgut does not harbor adult stem cells, further suggesting that adult ileal cells possess molecular mechanisms that enable their longevity (Sawyer, Cohen, and Fox 2017).

**Figure 1.**
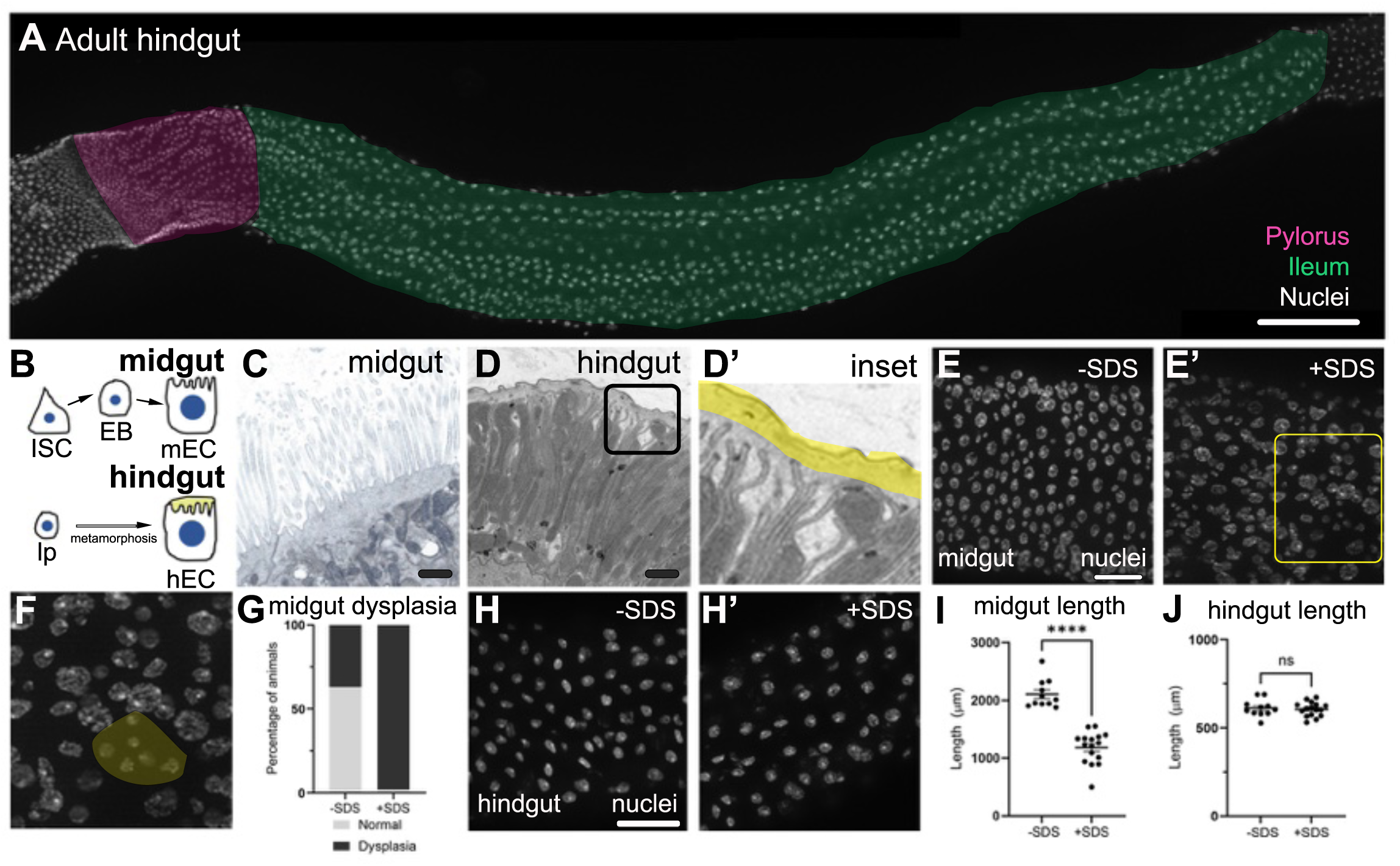
Adult hindgut enterocytes are long-lived and resist SDS. **A)** Image of the adult *Drosophila* hindgut. Pylorus and Ileum are pseudo-colored (midgut is to the left of the pylorus). White= nuclei, scale bar= 100µm. **B)** Illustration of the adult midgut EC lineage and larval pyloric cell to adult EC lineage. Intestinal stem cell (ISC), enteroblast (EB), midgut enterocyte (mEC), larval pyloric cell (lp), and adult hindgut enterocyte (hEC). Yellow shows cuticular layer. **C)** Electron micrograph of the apical (lumen facing) surface of an adult midgut EC, scale bar= 1µm **D)** Electron micrograph of the apical (lumen facing) surface of an adult hindgut EC, scale bar= 1µm. Black box highlights inset shown in **D’**. Yellow pseudocolor in **D’** shows cuticular layer. **E)** Adult midgut epithelium from an animal fed a control diet. White= nuclei, scale bar 25µm. **E’)** Adult midgut epithelium from an animal fed an SDS-containing diet. White= nuclei, scale bar= 25µm. Yellow box= highlights inset seen in **F. F)** Inset from yellow box in **E’**, highlighting a cluster of expanded small nuclei (non-ECs) in yellow. **G)** Normal vs. dysplastic adult midguts in animals fed control or SDS-containing diet. N= 67 -SDS and 49 + SDS animals, from 10 biological replicates. The difference in the distribution of gut morphologies between -SDS and +SDS is significant by Fisher’s exact test (p<0.00001) **H)** Adult ileal epithelium from an animal fed a control diet. White= nuclei, scale bar= 25µm. **H’)** Adult ileal epithelium from an animal fed an SDS-containing diet. White= nuclei. **I**) Length of the adult midgut in animals fed control or SDS-containing diet. N=11 no SDS, N= 16 SDS, from 3 biological replicates. Significant difference in midgut length between conditions by unpaired t-test (p<0.0001). **J)** Length of the adult hindgut in animals fed control or SDS-containing diet. N=11 no SDS, N= 16 SDS, from 3 biological replicates. No significant difference in hindgut length between conditions by unpaired t- test.

To identify cellular differences between adult midgut and hindgut ECs, we examined these ECs by electron microscopy. Both midgut and hindgut ECs contain extensive apical microvilli, which are typical of ECs (**Fig1C,D**). In the midgut, these microvilli are in direct contact with the contents of the intestinal lumen (**Fig1C**). However, in the hindgut, these microvilli are covered by a protective chitinous cuticular layer (**Fig1D,D’**). This cuticular layer is also seen in the ileum of ants (Villaro et al. 1999), the larval *Drosophila* ileum (Murakami and Shiotsuki 2001), and the adult *Drosophila* foregut (Zhu, Ludington, and Spradling 2024). This protective layer may extend the lifetime of adult hindgut ECs.

Given the longevity and protective chitinous structure of hindgut ileal ECs, we assessed whether these cells resist an ingested damaging agent that commonly disrupts the intestinal epithelium. Sodium Dodecyl Sulfate (SDS) is a detergent that induces EC loss in a variety of experimental models (Anderberg and Artursson 1993; Buchon, Broderick, Chakrabarti, et al. 2009; Welch et al. 2021; Ogulur et al. 2023). In *Drosophila,* SDS feeding induces adult midgut dysplasia, which is most apparent by the expansion of ISCs and their enteroblast daughters, both of which contain small diploid nuclei compared to the larger polyploid midgut EC nuclei (Marchetti, Zhang, and Edgar 2022). SDS feeding also decreases the length of the fly digestive tract, though whether this decrease specifically occurs in the midgut and/or hindgut was unclear (Pan and Jin 2014). We fed SDS to adults for two days (see methods) and assessed dysplasia and gut length in both the midgut and hindgut. SDS feeding induces midgut dysplasia, noted by the appearance of numerous clusters of small nuclei (**Fig1E,F,G**) and decreases midgut length (**Fig1I**). However, we observe no obvious alterations in hindgut EC nuclear morphology, or a change in hindgut length (**Fig1H,J**). Similarly, the adult hindgut pylorus remains unchanged in appearance upon SDS feeding (**SFig1A vs. B**). Previously, we found that adult ileal cells infrequently enter the cell cycle after day two of adulthood, suggesting that ileal cells do not turn over frequently (Fox and Spradling 2009). In further support, we find that the total cell number of the ileum does not change significantly between day two and thirty-five of adulthood (**SFig1C**). Taken together, our data highlight long-lived, protective, and stress-resistant properties of hindgut ECs, which distinguish them from midgut ECs.

### Adult hindgut ileum enterocytes are resistant to apoptotic stimuli upstream of caspase activation

Given the differences in cell turnover, SDS response, and ultrastructure between ECs of the hindgut and midgut, we next examined whether hindgut ECs are refractory to pro-apoptotic signaling. Pro-apoptotic genes at the H99 locus (*grim*, *reaper/rpr*, and *head involution defective/hid)* are targets of the transcription factor p53, and function to relieve inhibition of the *Drosophila* Inhibitor of Apoptosis 1 (Diap1). Diap1 inhibition then enables initiator caspase activity and apoptosis (Zhou et al. 1995; Zhou et al. 1997; White et al. 1994; Yoo et al. 2002; Muro et al. 2006; Grether et al. 1995; Chen et al. 1996; Brodsky et al. 2000; Chai et al. 2003; Olson et al. 2003) (**Fig2A**). Previous work in the midgut established that adult-onset overexpression of *reaper* induces EC apoptosis and midgut dysplasia (Jiang et al. 2009). Similarly, we find that adult-specific overexpression (see methods) of *UAS-hid* (*hid^OE^*) and *UAS-reaper (rpr^OE^)* with the midgut EC driver *Myo1A-Gal4* alters the regular pattern of nuclei and EC nuclear size (**Fig2B,B’**), a sign of apoptosis-related midgut dysplasia (Jiang et al. 2009). In contrast, we find the expression of *UAS-dronc* (*dronc^OE^*) with *Myo1A-Gal4* does not appreciably disturb midgut morphology (**Fig2B”**). Dronc overexpression alone is typically not a potent driver of cell death, as feedback mechanisms within a Dronc-containing complex known as the apoptosome control the degree of apoptosis (Shapiro et al. 2008; Florentin and Arama 2012). Further, unlike *hid* or *rpr, dronc* is known to be expressed without causing apoptosis in the adult midgut, where it functions in preserving the intestinal barrier (Lindblad et al. 2021).

**Figure 2.**
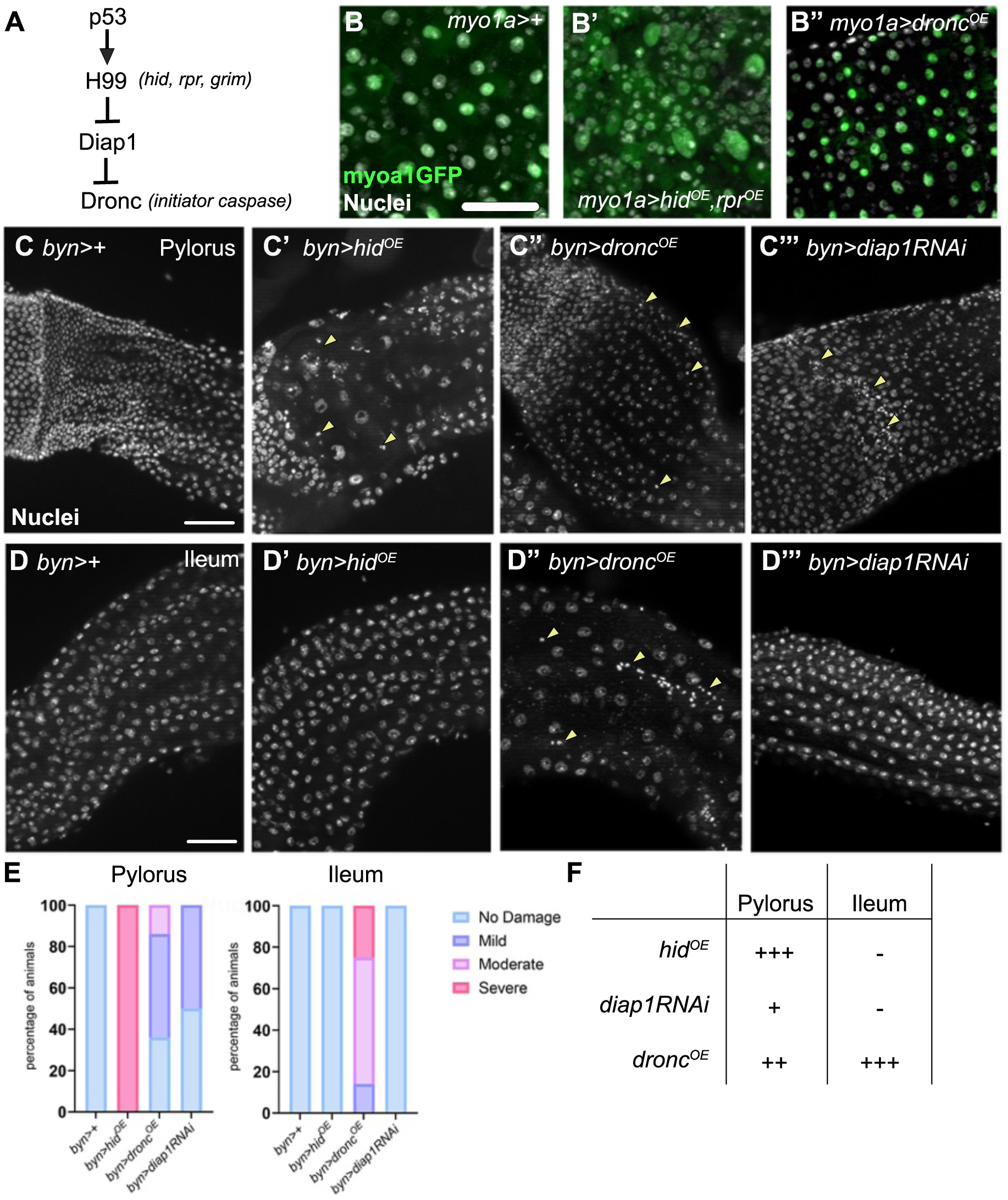
Upstream caspase signaling cannot initiate cell death in the adult ileum. **A)** Simplified diagram of the signaling cascade that controls *Drosophila* initiator caspase activation. **B)** Adult midgut enterocytes expressing *myo1a Gal4>UAS-GFP* **(B)**, *myo1a Gal4>UASGFP, UAShid, UASrpr (=hid^OE^, rpr^OE^)* **(B’)** or *myo1a Gal4>UASdronc (=dronc^OE^)* **(B’’)**. White= nuclei, Green=GFP, scale bar= 50µm. N= 6, 2 biological replicates. **C)** Adult pylorus. Wild type, 4d, N=29 **(C)**, expressing *byn Gal4> UAS-hid,* 4d, N=25 **(C’)**, expressing *byn Gal4> UAS-dronc,* 4d, N=22 **(C”)**, expressing *byn Gal4> UAS-diap1RNAi,* 10d, N=10 **(C’’’)** White= nuclei, yellow arrowheads= pyknotic nuclei, scale bar= 50µm. 3-6 biological replicates. **D)** Adult ileum. Wild type, 4d, N=29 **(D)**, expressing *byn Gal4> UAS-hid,* 4d, N=25 **(D’)**, expressing *byn Gal4> UAS-dronc,* 4d, N=22 **(D”)**, expressing *byn Gal4> UAS-diap1RNAi,* 4d, N=10 (**D’’’)**. White= nuclei, yellow arrowheads= pyknotic nuclei, scale bar= 50µm. 3-6 biological replicates. **E)** Quantification of the extent of damage in the pylorus and ileum, see methods. In control conditions there is no significant difference between the pylorus and ileum (Fisher’s exact test). In the *hid^OE^*, *dronc^OE^*, *diap1RNAi* conditions, there is a significant difference between the pylorus and ileum (Fisher’s exact test, p<0.0001). **F)** Table summarizing the relative strength of cell killing by the indicated transgene in the adult pylorus and ileum.

Next, we examined the sensitivity of midgut progenitor cells (ISCs and EBs) to cell pro-apoptotic signals. Using the midgut progenitor driver *esg-Gal4,* we observe almost complete loss of GFP-labeled adult midgut progenitors upon *hid* induction (**FigS2A,A’**). In contrast, induction of *dronc* in midgut progenitors does not lead to appreciable cell loss (**FigS2A’’**). Again, this result is consistent with the idea that *dronc* is often not sufficient to induce apoptosis. Moreover, *dronc* expression functions to preserve the adult midgut progenitor pool in a non-apoptotic role (Arthurton et al. 2020). To summarize our findings in the adult midgut, we find that activating signals upstream of caspases, but not the *dronc* caspase itself, severely disrupts the midgut epithelium.

We next compared our midgut findings to that of the hindgut. We previously showed that adult-specific expression (see methods) of either *rpr* or *hid/rpr* together, using the pan-hindgut driver *brachyenteron*- (*byn)-Gal4* leads to pyknotic nuclei and cell loss in the pylorus (Fox and Spradling 2009; Sawyer, Cohen, and Fox 2017; Cohen et al. 2018). We reproduced these results using *byn-Gal4* to drive *hid* expression in the adult hindgut. In the adult pylorus, as expected, we observe *hid-*induced cell loss and nuclear hypertrophy (larger nuclei), which is a known pyloric regenerative response (Losick, Fox, and Spradling 2013; Sawyer, Cohen, and Fox 2017; Cohen et al. 2018; Cohen et al. 2021) (**Fig2C’**). We further confirmed the ability of *hid* to induce pyloric apoptosis using *CD8::PARP::VenusGFP* (*CPV*), an artificial effector caspase substrate (Florentin and Arama 2012). In this construct, a CD8 membrane tag is connected to an intracellular VenusGFP tag by a linker that contains a caspase recognition sequence. Caspase cleavage exposes a PARP antibody epitope, which indicates caspase activity. In the adult pylorus, *hid* induction leads to robust cleaved substrate recognized by the PARP antibody (**FigS2D,E,F**). Next, we examined *dronc* overexpression in the pylorus. In comparison to wildtype animals, we see mild to moderate damage in the pylorus, with single or small groups of pyknotic nuclei (**Fig2C”,E**). While this level of apoptosis upon *dronc* in the pylorus is greater than that of what we find in the midgut, overall, our results are consistent in that H99 genes are more potent than *dronc* in promoting cell loss in the midgut ECs, the midgut progenitors, and in the hindgut pylorus.

In contrast to the above findings in the midgut and hindgut pylorus, our results with *hid* and *dronc* expression in the ileum are completely different. We observe no change in the nuclear appearance of ileal ECs upon *hid* expression, indicating that these ECs are refractory to *hid-*induced apoptosis (**Fig2C’vsD’**). Consistent with this finding, in animals that co-express *hid* and *CD8::PARP::VenusGFP* (*CPV*), PARP antibody does not accumulate in the adult ileum (**SFig2D’, E’, F**). In further contrast to the pylorus, *dronc* overexpression in the ileum leads to appreciable cell loss with large patches of pyknotic nuclei (**Fig2D”,E**). Of note, the ability of *dronc* expression to ablate adult ileal ECs rules out the possibility that the *byn-Gal4* driver is not appreciably expressed in the adult ileum, as does our observed expression of VenusGFP from the CPV transgene (**FigS2E’**). Next, to assess if our hindgut expression system induces cell ablation via apoptosis, we expressed either the baculovirus protein p35, known to prevent apoptotic death (Hay, Wolff, and Rubin 1994), or the caspase inhibitor *diap1.* We find the overexpression of *p35* or *diap1* rescues *hid* damage in the pylorus and *dronc* damage in the pylorus and ileum, consistent with the idea that the cell death events that we observe are due to apoptosis (**SFig2G**). Together, our results suggest that adult ileum ECs are unique due to their insensitivity to H99 gene expression and sensitivity to *dronc* expression (**Fig2F**).

As H99 expression relieves Diap1 inhibition of the caspase Dronc (**Fig2A**), we next examined the impact of reducing Diap1 in mature adult hindguts. We knocked down *diap1* by RNAi using *byn-Gal4.* After 4d of *diap1RNAi,* we see no appreciable cell death in the adult pylorus or ileum (**FigS2B-C**). *diap1RNAi* expression in the hindgut during embryogenesis and larval development leads to organismal death, while animals with *diap1RNAi* induced during early pupal development die soon after eclosion and lack hindguts, demonstrating that embryonic and larval hindgut cells are sensitive to *diap*1 loss (data not shown). Most likely, while 4d induction is sufficient for us to observe caspase signaling gain of function phenotypes (*hid^OE^, dronc^OE^),* the 4d RNAi is insufficient to deplete all endogenous Diap1. Therefore, we extended the induction of *diap1RNAi* to 10d in adult hindguts. After this longer knockdown period, we indeed see pyknotic nuclei in the pylorus, but not the ileum (**Fig2C’’’, D’’’**). These results suggest that adult ileum ECs are more resistant to *diap1* loss than pyloric cells (**Fig2F**), which is consistent with the idea that the ileum is refractory to apoptotic regulation upstream of *dronc*. Together, these findings indicate that long-lived, SDS-resistant ECs of the adult ileum are uniquely wired for caspase signaling, and in particular are highly refractory to *hid* induced apoptosis (**Fig2F**).

### Ileum EC ablation by the caspase Dronc does not elicit a robust regenerative response

The ability of *dronc* activation to ablate adult ileal ECs enabled us to examine whether the adult ileal ECs can regenerate. Ablation of either midgut ECs or midgut progenitors in adults elicits a regenerative response, most notably by increased ISC division and subsequent EC differentiation (Apidianakis et al. 2009; Buchon, Broderick, Poidevin, et al. 2009; O’Brien et al. 2011; Amcheslavsky, Jiang, and Ip 2009; Jiang et al. 2009; Jiang et al. 2011; Staley and Irvine 2010; Zhai, Boquete, and Lemaitre 2017). Additionally, both adult midgut ECs and hindgut pyloric cells are capable of injury-induced endocycles, which regenerate lost epithelial mass by producing large polyploid cells (Xiang et al. 2017; Losick, Fox, and Spradling 2013; Cohen et al. 2021; Cohen et al. 2018). If the ileum is regenerated, this might occur through endocycles of remaining ileal cells, as occurs in the pylorus. Alternatively, pyloric cells could produce new ileal cells, as occurs during developmental adult hindgut formation (Fox and Spradling 2009; Takashima et al. 2008; Zhang et al. 2024).

To test whether either ileum EC ablation leads to ileum regeneration, we expressed *dronc* in adult hindguts for 4d as before and then allowed animals to recover for 6d. After 4d of *dronc* expression, we observe some cell loss and pyknotic nuclei compared to control animals in the ileum (**Fig3A vs. 3A’,B, 2D”**). However, after 6d of recovery we observe extensive cell loss in the ileum and large patches pyknotic nuclei (**Fig3A vs. 3A”,B**). Some ileal cells appear larger, suggesting some may increase their ploidy or undergo nuclear decondensation. However, this response is quite weak compared to the polyploid regeneration in the pylorus after *hid* induction (**Fig2C’**). Upon *dronc* induction in the pylorus followed by a 6d recovery, we observe pyknotic nuclei and some cell loss, though not as extensive as seen in the *dronc-*expressing ileum or *hid*- expressing pylorus (**FigS3A-B, Fig2C’’**). In summary, we find that the long-lived ECs of the adult hindgut ileum resist most cell death stimuli but are acutely sensitive to direct activation of the Dronc caspase. Upon Dronc activation, ileal cells are depleted, but are not readily replaced through a regenerative mechanism.

**Figure 3.**
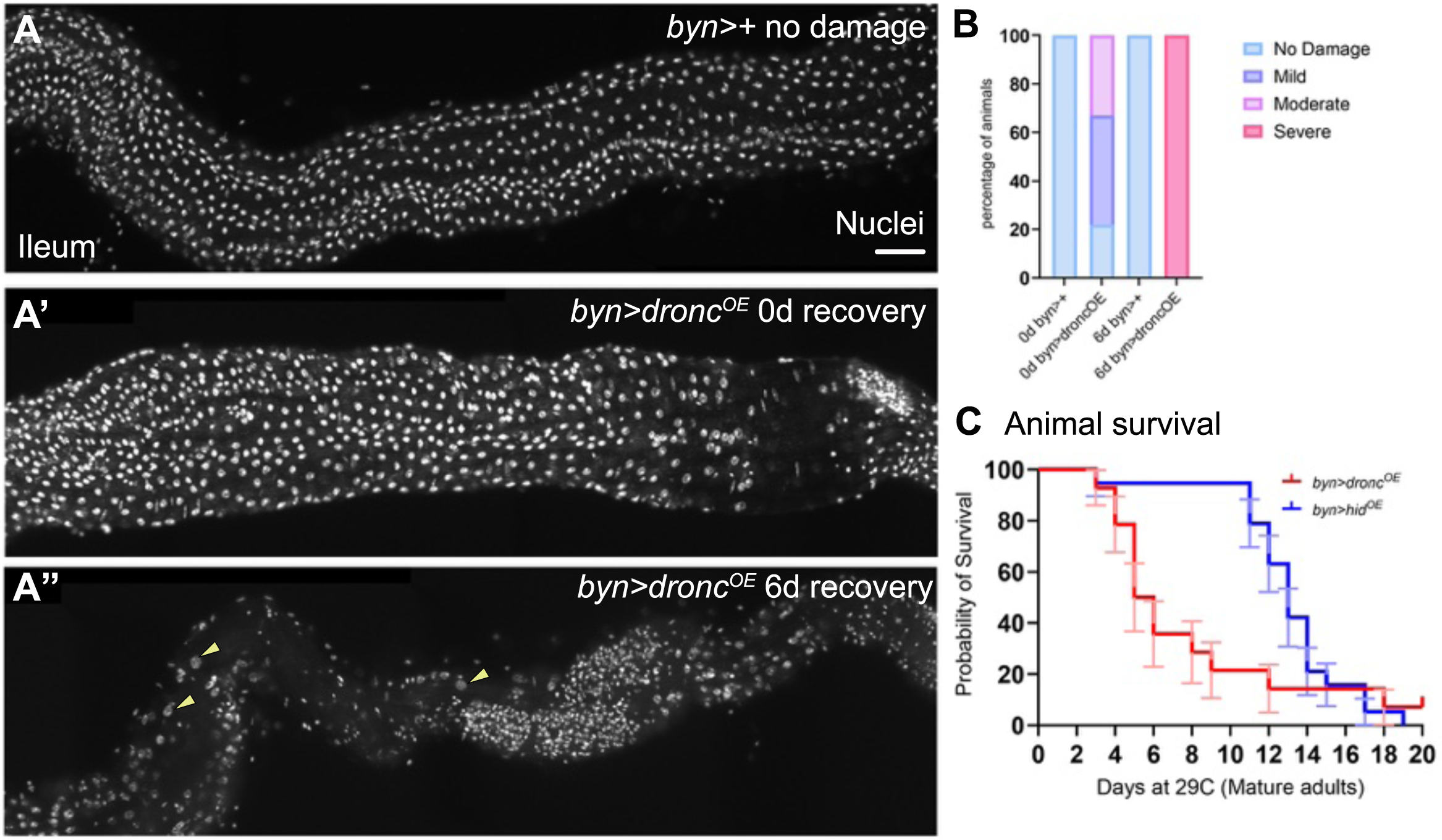
Dronc overexpression causes ileal EC death, but not a robust regenerative response. A-A’’) Adult ileum. White=nuclei, yellow arrowheads= large ileal nuclei, scale bar= 50µm. **A)** Ileum, no *dronc^OE^* damage, N=24. **A’)** Ileum, 4d *dronc^OE^*, 0d recovery, N=9. **A”)** Ileum, 4d *dronc^OE^*, 6d recovery, N=17. **B)** Quantification of the extent of ileal damage, 3-4 biological replicates, Fisher’s exact test, p<0.0001. **C)** Survival curve of adult *hid^OE^* (N=19) vs. *dronc^OE^* (N=14) animals. 2 biological replicates.

Our ability to ablate adult hindgut ileal ECs, coupled with the inability of this tissue to regenerate, enabled us to examine the consequences of ileal cell depletion. Within 5d of continuous *dronc* expression, roughly 50% of animals die (**Fig3C**). By comparison, it takes roughly two weeks to reach 50% lethality in continuous *hid-*expressing animals, which have a damaged and actively regenerating pylorus but fully intact ileum (**Fig3C**). This difference likely reflects both the larger area covered by the ileum compared to the pylorus, as well as the lack of a regenerative mechanism to combat injury. These results highlight the importance of ileal tissue integrity for animal survival and highlight the acute sensitivity of adult *Drosophila* to loss of this long-lived barrier epithelium.

### Ileum cell death resistance arises during adult gut maturation and coincides with substantial transcriptome changes

Our data highlight differences between hindgut ileum ECs and other gut epithelial cells in terms of sensitivity to apoptotic gene expression and in regeneration. Within the hindgut, the differences between the pylorus and ileum are particularly intriguing, given that these cell populations arise from a common progenitor pool during metamorphosis (Fox and Spradling 2009; Takashima et al. 2008; Zhang et al. 2024). We therefore leveraged the predictability of gut developmental timing to pinpoint when the pylorus and ileum diverge in terms of resistance to *hid* cell killing. Unlike in mature adults, we observe that the ileum of newly eclosed (≤24h) adult flies is sensitive to *hid*-induced cell death, as indicated by robust loss of nuclei upon *hid* expression as is seen in the pylorus (**Fig4A, A’, B, B’**). The ileum of young *hid*-expressing animals is also highly active in terms of the cell cycle, as indicated by BrdU incorporation, similar to that of the *hid* pylorus (**Fig4B’,E,F**). This BrdU activity likely reflects a regenerative capacity of the young (1-2 day old) ileum. These results are in striking contrast to the lack of cell loss and BrdU incorporation in the mature adult ileum of *hid* expressing animals (**Fig4D- D’,F**). In contrast to the mature ileum, the mature pylorus remains sensitive to *hid* expression, indicated by the loss of pyloric cells and the incorporation of BrdU upon *hid* expression (**Fig4C-C’, E**). Therefore, the point of divergence in *hid* cell death resistance between the adult pyloric and ileal epithelia occurs roughly 24h after eclosion.

**Figure 4.**
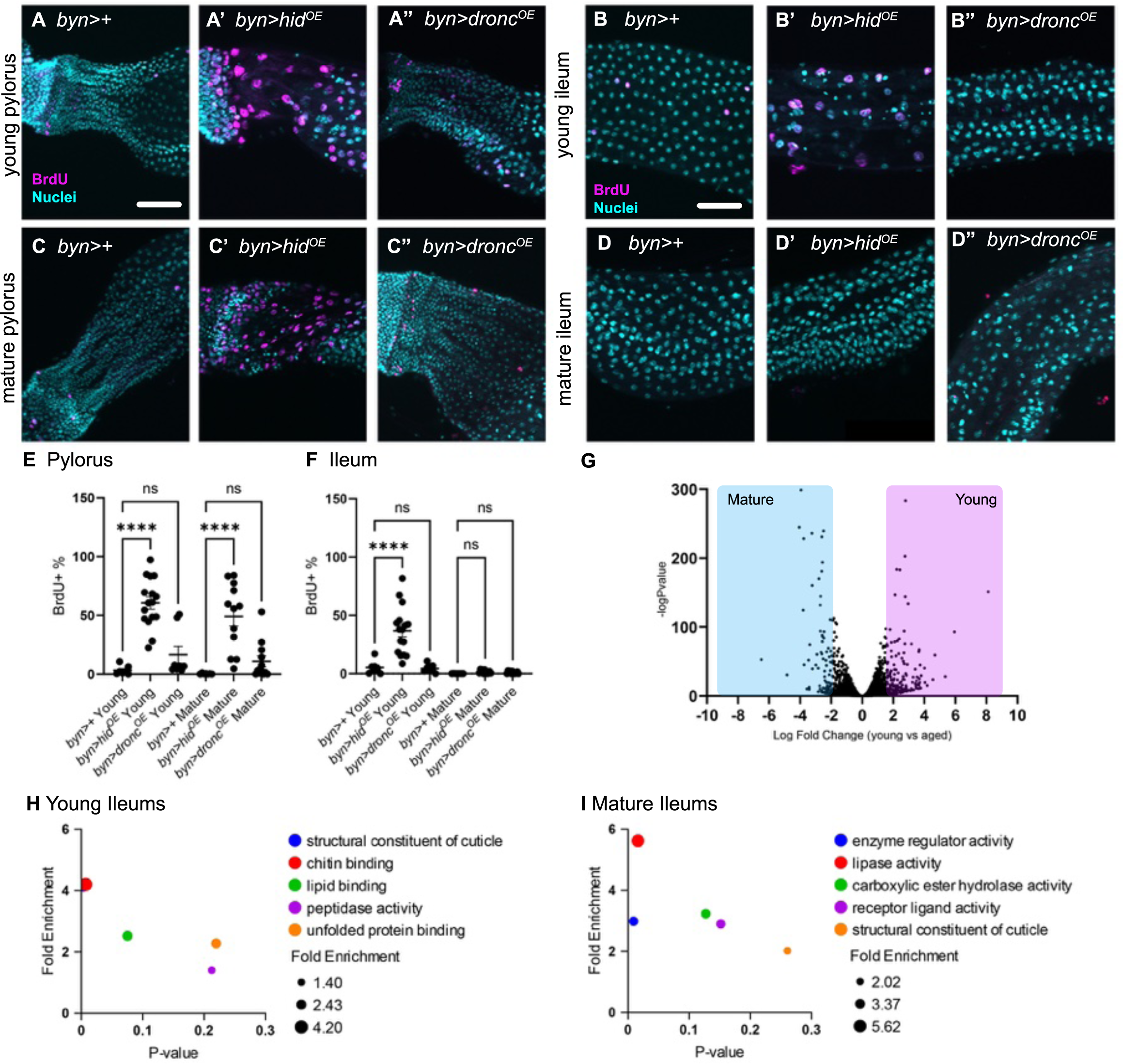
Mature and young ileums have different responses to injury and distinct transcriptional profiles. A-D’’) Adult hindguts. Magenta= BrdU, Cyan= nuclei, scale bar= 50µm. Genotypes noted in figure. **A-B’’)** S-phase activity in *hid^OE^* or *dronc^OE^* young adult hindgut pylorus and ileum, less than 24hrs old. **C-D’’)** S-phase activity in *hid^OE^* or *dronc^OE^* mature adult hindgut pylorus and ileum, aged for 4d at 18C. **E-F)** Percentage of BrdU+ cells in animals of the indicated ages and genotypes. Each data point equals one animal. 3 biological replicates. **G)** Volcano plot representing RNAseq data comparing young and mature ileums. **H-I)** Pangea Reactome Enrichment Analysis of genes with a greater than 2-log fold change.

Interestingly, we see very little BrdU incorporation after *dronc* expression in the pylorus or ileum of young or mature animals (**Fig4A”,B”,C”,D”,E, F**), suggesting that *hid* expression may induce a more robust regenerative response than *dronc* expression.

Given the long-lived nature of ileal ECs, we examined candidate markers that might be enhanced in a long-lived cell population. As longer-lived cells may acquire DNA damage, we examined the accumulation of DNA damage foci over time, using the marker γH2Av. Indeed, adult ileal ECs exhibit an increase in γH2Av foci when comparing two-day vs. five-day old animals (**SFig4A-A’ vs. C-C’,D**). Jun Kinase signaling is linked to cell survival, notably under stress conditions (Cosolo et al. 2019). Therefore, we examined a marker of JNK signaling, whereby the JNK-responsive tetradecanoylphorbol acetate response element (TRE) is fused to dsRed (Chatterjee and Bohmann 2012). As for γH2Av, the percentage of TRE-DsRed cells in the ileum increases when comparing two-day vs. five-day old animals (**SFig4A,A’’ vs. B-B’,D**). These markers highlight differences in stress-associated gene expression in adult hindgut ECs.

To further uncover important gene expression changes that underlie cell preservation in the ileum, we conducted RNAseq of ≤24h (=young) and 96h (=mature) adult hindguts (methods, **Fig4G**). We analyzed the Gene Ontology (GO) terms of genes with a ≥2-fold change between young (N=91) and mature (N=132, **Suppl. Table 1**). In young animals, we find an enrichment of chitin and cuticle associated genes (**Fig4H**). Our analysis fits with the idea that the protective chitinous, cuticular layer seen in the adult ileum (**Fig1D- D’**) is built in early adulthood. In mature animals, we see an enrichment in enzymatic and lipase genes (**Fig4I**), which fits with the role of the mature adult ileum in resorption (Cohen et al. 2020). Overall, our findings highlight important differences between the young and mature ileum in terms of caspase signaling, cell cycling, and gene expression.

### The cell adhesion protein *CG15312/Loofah* promotes ileum EC cell death resistance

Our RNAseq uncovered numerous changes that may regulate cell death resistance in the ileum. To identify such regulators, we conducted a candidate RNAi screen of 82 genes that are elevated in mature ileums. We conducted this screen in a *hid-* overexpression background, to identify those regulators that preserve the ileal epithelium in response to this apoptotic stimulus **(Fig5A**). To identify genes that are critical for ileal function, we used animal survival as a readout in a primary screen. As a positive control, we first examined animals co-expressing *hid* and *diap1RNAi.* After ten days of transgene co-expression, we observe ∼75% animal lethality (**Fig5C**). Expression of either transgene alone causes little lethality after 10 days, highlighting the synergy of co-expression of these two death-promoting transgenes **(Fig5C**). We reasoned that genes which are upregulated in mature hindguts may counteract *hid*- induced cell death, and therefore RNAi of such genes in *hid* expressing animals will result in lethality within ten days, similar to the synergistic cell death seen in *hid* and *diap1RNAi* animals.

**Figure 5.**
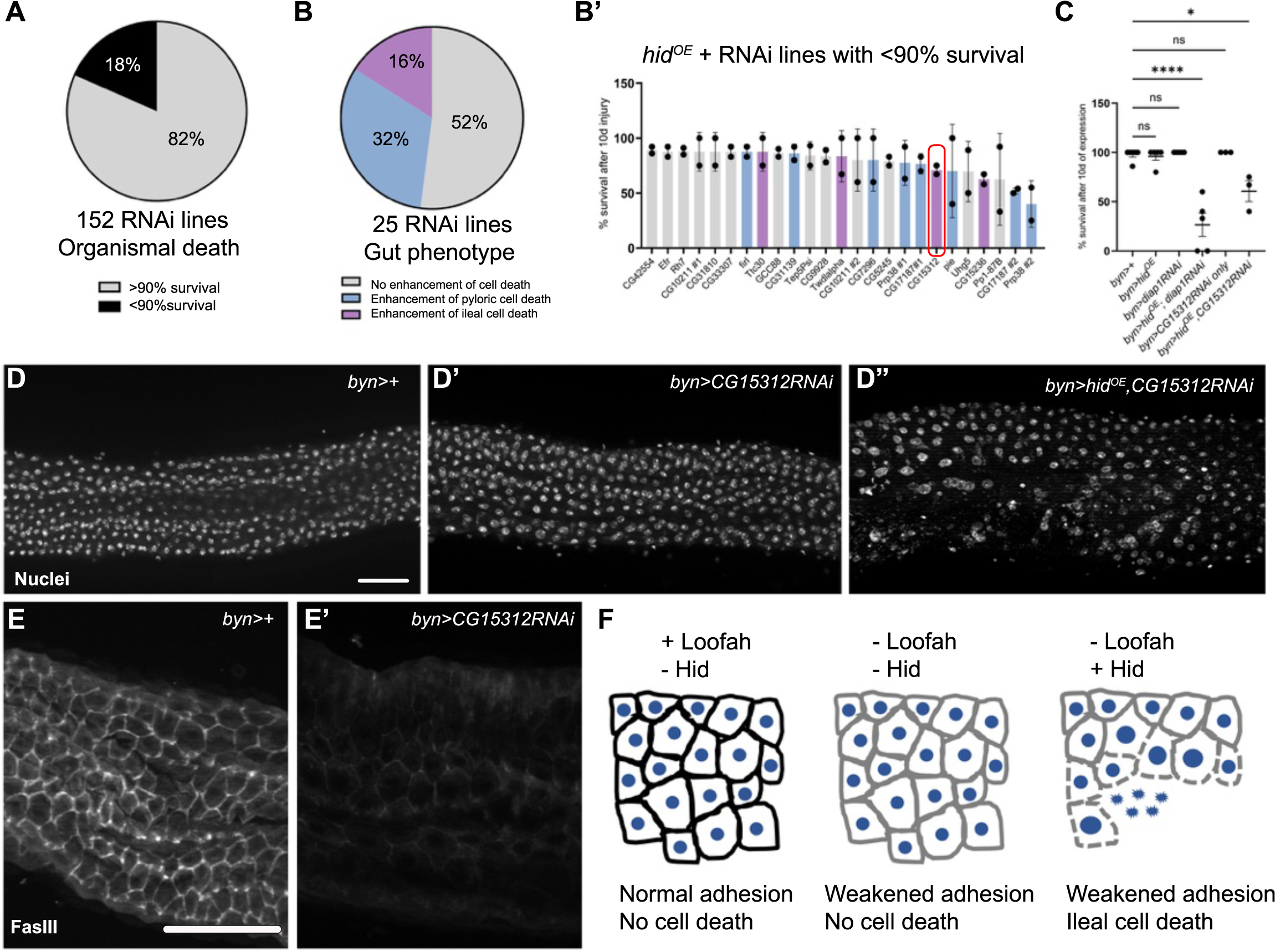
C*G15312/*Loofah, an immunoglobulin superfamily cell adhesion protein, prevents ileal cell death in response to *hid* expression. A) Outcome distribution from a primary screen of 152 RNAi lines of 82 genes identified by RNAseq. Each transgene was expressed for ten days, after which survival was measured. **B)** Outcome distribution of Ileal phenotypes from the secondary imaging-based screen. **B’)** Individual line survival data from the primary screen of the animals of the indicated phenotypes. Colors for each bar indicate the category of each line in the secondary imaging-based screen in **B**. Each data point equals one animal. 2 biological replicates. *CG15312* is highlighted in a red box. **C)**. Plot comparing survival after 10d of the indicated phenotypes. 2-3 biological replicates. ANOVA, Tukey’s test, p<0.0001. **D)** Adult ileums after 10d of transgene expression, genotypes indicated. White= nuclei, scale bar= 50µm. **E-E’)** Comparison of FasIII localization after ten days of transgene expression, genotypes indicated. White= FasIII, scale bar= 50µm. **F)** Model of Loofah function, and the impact of *hid* expression, in the adult hindgut.

Our primary organismal lethality screen consisted of 152 RNAi lines, chosen from the top 82 upregulated genes in mature ileums, including two RNAi lines per gene when possible. From this initial screen, 28 lines showed ≤90% survival and were selected for further screening (**Fig5A, B’, Suppl. Table 2**). We conducted a secondary microscopy-based screen to assess the hindgut phenotype of 25 of these lines (**Fig5B**). We identified three classes of phenotypes: no enhanced cell death (13 lines), enhanced pyloric cell death (8 lines), and enhanced ileal cell death (4 lines, **Fig5B**). Genes found in our screen that enhanced ileal cell death were *Ttc30, Twdlalpha, CG15236, and CG15312.* Thus, our RNAseq analysis revealed multiple gene expression changes that promote ileal cell survival upon *hid* expression.

We further characterized *CG15312,* as RNAi of this gene leads to significant ileal cell loss upon *hid* induction (**Fig5D**). Further, *CG15312* has clear homology to cell adhesion proteins in several organisms, including immunoglobulin superfamily proteins in humans, rodents, frogs, and nematodes ((Bult and Sternberg 2023)**SFig5A**). Only one RNAi line for *CG15312* is available, therefore we confirmed knockdown of the gene with RT-qPCR (**SFig5B**). Loss of *CG15312* gene function alone via RNAi for ten days in the adult hindgut does not result in organismal cell death or appreciable changes in hindgut nuclear morphology (**Fig5C,D’**). In contrast, loss of *CG15312* in a *hid* overexpression background leads to reduced animal viability (**Fig5B’, C**) and significant ileal cell loss (**Fig5D”**). We observe differences in nuclear morphology, with some remaining nuclei appearing larger and patches of pyknotic nuclei (**Fig5D”**). These clusters resemble cellular phenotypes seen in many cell adhesion/polarity mutants (Muller 2000). To test whether *CG15312* impacts ileal cell adhesion in the absence of *hid* expression, we examined the localization of FasIII, which prominently localizes to ileal cell junctions (Sawyer, Cohen, and Fox 2017; Takashima et al. 2008). Loss of *CG15312* decreases FasIII localization at ileal cell junctions, consistent with a role in cell adhesion (**Fig5E**). Our data suggests that loss of *CG15312* in *hid-*expressing ileal cells leads to loss of cell-cell adhesion and cell death. Given our findings, and the lack of a previous characterization of *CG15312,* we name this gene product *<u>lo</u>w <u>o</u>n membrane <u>fa</u>s and enhancer of <u>h</u>id* (*loofah*), due to the loss of a cell membrane component and subsequent cell loss that leads to a disorganized epithelium. In summary, hindgut ileum maturation involves upregulation of the conserved yet uncharacterized cell adhesion gene *loofah. loofah* expression promotes stability of the adhesion protein FasIII on ileal EC membranes and promotes cellular longevity through resistance to death signaling upstream of caspases.

## Discussion

To achieve regenerative tissue therapy, we must not only understand how to regrow tissue lost to injury but also must uncover ways that tissue can be maintained and withstand insults. Currently, regenerative research heavily favors study of regrowth. In this study, we present a new tissue model, the hindgut ileum, as an accessible system to study tissue preservation. In doing so, we reveal a role for a completely uncharacterized yet conserved cell adhesion protein, which we name Loofah, in repression of cell death.

### Ileal cells are a model of an epithelium that is built to last

Unlike most intestinal cell types, ileal ECs are resistant to *hid*-induced cell death. Our data suggests that ileal ECs utilize multiple mechanisms to survive long-term. First, the thick extracellular cuticle may protect ileal ECs from ingested toxins. Second, resistance to upstream caspase signaling is a mechanism of ileal cell preservation. As we show here, three different intestinal cell types- the midgut progenitors, the midgut ECs, and the hindgut pyloric cells- are all sensitive to *hid* expression, whereas the ileum is not. Our data suggest that a combination of structural features and apoptosis resistance enables ileal cells to persist throughout the lifespan of the organism. Avoiding cell death is an important strategy for survival of ileal ECS, as they have a limited regenerative capacity.

Our data here and that of others highlight the complexity of caspase signal regulation. Clearly, the caspase activation cascade is not a simple linear pathway, but rather each step in the pathway can have different degrees of regulation. One example of this intricacy in caspase regulation from our study here is that ileal ECs are acutely sensitive to overexpression of Dronc, whereas pyloric hindgut cells are not. This finding suggests that ileal cells may primarily suppress apoptosis through repression of initiator caspases. When Dronc is ectopically activated, this strong repression is bypassed. In support of this finding, our results highlight that ileal cells are not as sensitive as pyloric cells to loss of the Dronc inhibitor Diap1. Many previous studies of the *Drosophila* caspase signaling cascade have revealed layers of feedback within the pathway, and the importance of the levels of each pathway component (Herriage, Huang, and Calvi 2024; Sulekh et al. 2024; Ziraldo and Ma 2015; Waldhuber, Emoto, and Petritsch 2005; Quinn et al. 2000; Wells, Yoshida, and Johnston 2006). The ileum may lack some of these feedback nodes, or may integrate feedback in a unique manner such that *hid* activation is unable to promote cell death. Future studies can identify the apoptotic network rewiring that promotes ileal EC longevity, which may ultimately provide unique targets for therapeutic tissue preservation. Further, it would be interesting to determine if this resistance extends to other forms of cell death, such as autophagy, necrosis, and ferroptosis. Alternatively, ileal ECs may maintain low levels of initiator caspases for non-apoptotic functions. If caspase levels are low, initiation of *hid* may not be sufficient to induce cell death. Indeed, caspases play roles in tissue integrity and differentiation (Gorelick-Ashkenazi et al. 2018; Fernando et al. 2002; Weaver et al. 2020; Fan et al. 2014; Klemm, Van Hazel, and Harris 2025; Nair and Baker 2024; Rosa et al. 2024; Galasso et al. 2023). In addition, the unique sensitivity of the ileum to *dronc* expression, and the rapid death of flies expressing *dronc,* may be of interest in efforts for targeted insect population control.

Regional cellular diversity along the anterior/posterior axis is a conserved property of the metazoan intestinal tract (Zwick et al. 2024; Marianes and Spradling 2013). Study of regional differences in the intestine has revealed fascinating biology, such as distinct commensal niches (Zhu, Ludington, and Spradling 2024), cancer vulnerabilities (Jiang et al. 2017), and metabolic programs (Zwick et al. 2024). Our data further highlights the importance of understanding the biology of diverse cell types to reveal new tissue maintenance strategies.

### Loofah, a novel regulator of cell death signaling

In addition to establishing the ileum as a new model of cellular longevity, our study reports a new role for an uncharacterized gene, *CG15312/loofah.* Our *hid*-enhancer screen reveals Loofah as a novel adhesion gene that suppresses cell death in ileal ECs. Loofah acts to preserve FasIII, a component of the septate junction cell adhesion complex. Previous work has highlighted some connections between cell adhesion and cell death. For example, loss of integrin adhesion can lead to autophagy (Vlahakis and Debnath 2017). Further, Protogenin, a mammalian ortholog of Loofah, promotes integrin adhesion which in turn suppresses apoptosis in the developing mouse skull (Wang et al. 2013). In our work, loss of Loofah in combination with *hid* overexpression has similarities to a form of cell death known as anoikis, where cell death occurs through loss of contact with the extracellular matrix(Taddei et al. 2012). Resistance to anoikis is a hallmark of cancer progression (Paoli, Giannoni, and Chiarugi 2013). We speculate that a cell death signal, plus weakening of adhesion together is sufficient to trigger cell death in ileal ECs (**Fig5F**). It is tempting to speculate that Loofah may be involved in maintaining ileal enterocyte connections to the basement membrane, and that loss of this connection leads to cell death. Unlike the adult midgut, the ileal enterocytes may be limited in their ability to repair these connections (Stricker, Hutson, and Page-McCaw 2025), and this places greater importance on cell preservation mechanisms.

Our results demonstrate that the *Drosophila* hindgut ileum is a powerful model to understand the mechanisms by which cells can resist cell death and extend their longevity. Our work here highlights the potential of studying cellular longevity as a complementary approach to study of regeneration, and that such study may prove especially important in tissues with limited regenerative potential.

## Materials and methods

### Drosophila genetics and culturing

All flies were raised at 25°C on standard media unless noted otherwise (Archon Scientific, Durham NC). Fly strains used listed in **Supplemental Table 3**. Flybase (flybase.org) lists complete genotypes. For all adult cell death transgene experiments, the genotype included a temperature-sensitive Gal80 transgene driven by a Tubulin promoter (Gal80ts), as well as the indicated Gal4 and UAS-transgene. Flies with Gal80^ts^, Gal4, and a UAS-transgene were kept at 18°C until adulthood and shifted to 29°C at the indicated time of adulthood. For experiments involving a 6d recovery from injury, animals were shifted back to 18°C. We aged newly eclosed flies at 18°C for 4d, shifted to 29°C for 48hrs or 96hrs to induce injury. We dissected females for all experiments, unless otherwise noted. For the *hid*-enhancer screen, adults of the selected genotypes were raised at 18°C for 4d and then shifted to 29°C for ten days. Animal lethality was counted for 2 replicates. Lines that had less than 90% survival were repeated and females were dissected to examine the ileal phenotype.

### Tissue fixation, staining, and imaging-

Electron microscopy was performed as described in (Schoenfelder et al., 2014). For antibody staining, tissues were dissected in 1X PBS and immediately fixed in 1XPBS, 3.7% paraformaldehyde, 0.3% Triton-X for 15-45min depending on antibodies. Immunostaining was performed in 1XPBS, 0.3% Triton-X, and 1% normal goat serum. The following antibodies were used in this study: FasIII (DSHB, 7G10, Mouse, 1:50), BrdU (Serotec, 3J9, 1:200), γH2Av (DSHB, UNC93-5.2.1.s, Mouse 1:1000), PARP (Abcam, ab2317, Rabbit, 1:200), Hoechst 33342 stain was used to label DNA (Life Technologies, 1:1000). All secondary antibodies used were Alexa Fluor dyes (Invitrogen, 1:500). Tissues were mounted in Vectashield (Vector Laboratories Inc.). Images were acquired with the following: an upright Zeiss AxioImager M.2 with Apotome processing (10X NA 0.3 EC Plan-Neofluar air lens or 20X NA 0.5 EC Plan-Neofluar air lens). Image analysis was performed using ImageJ (Schneider et al., 2012), including adjusting brightness/contrast, image stitching, Z projections, and cell counts. Cell counts were acquired by using (1) the cell counter or (2) threshold function to identify objects. For cell death quantification, mild = some pyknotic nuclei, moderate= pyknotic nuclei and small patches of cell loss, severe = pyknotic nuclei and large patches of cell loss.

### SDS and BrdU Feeding assays

For SDS feedings, all animals were w1118. Females aged 4-5 days at 25°C were fed either 5% sucrose (EMD Chemicals) or 5% sucrose + 0.6% SDS (Sigma). For BrdU feedings, flies were fed 2.5mg/mL BrdU in 5% sucrose for 24hrs on day 2 of the 29°C shift. For midgut dysplasia quantification, midguts were examined for three or more discrete clusters of greater than three small progenitor-like cells without intervening ECs. Guts with such clusters were scored as dysplastic.

### RNA seq-

Female flies less than 24hrs old at 25°C and aged for 96hrs at18°C were collected for dissection. Only ileums were dissected from female animals. Guts were dissected in groups of 10 and then placed into cold Trizol (Life Technologies). ∼100-150 animals were dissected per condition for 3 replicates. RNA extraction protocols performed as previously described (Stewart et al. 2024). RNA samples were sent to Novogene (Beijing, China) for library generation and sequencing on an Illumina Novaseq machine to generate paired end, 150 base pair reads. The FASTQ data were first processed with TrimGalore! to remove adapters and low-quality reads. HISAT2 was used to align and mapped to the DM.BDGP6.32.105 genome downloaded from Ensembl. The feature counts function was used to count the reads per feature, then followed by DESeq2 to determine differential gene expression. We analyzed the Gene Ontology (GO) terms using the Reactome in Pangea. We used Slim2 GO Molecular Function enrichment for all genes with over a 2-fold difference (Hu et al. 2023).

### RT-qPCR-

Knockdown of Loofah was validated using RT-qPCR. We expressed Loofah RNAi (VDRC_KK) in salivary glands using the NP5169-Gal4 (NPGal4) driver at 29 °C. We used an RNAi control line (VDRC_60000) for comparison. Salivary glands were dissected on day 6 from wandering L3 larvae in ice-cold Schneider’s medium supplemented with 10% FBS. RNA extraction was performed as previously described (Stewart et al. 2024). Equal amounts of RNA were used for cDNA synthesis using the iScript™ cDNA Synthesis Kit (Bio-Rad, #1708891). A Luna® Universal qPCR Master Mix (NEB, #M3003L) was used for the RT-qPCR. We used amplification of *rp49* as the control (Janssens et al. 2017).

The RT-qPCR primers used to amplify *rp49* are as follows (164bp amplicon):

Forward: 5’-ATCGGTTACGGATCGAACAA-3’

Reverse: 5’-GACAATCTCCTTGCGCTTCT-3’

The RT-qPCR primers used to amplify *loofah* are as follows (101bp amplicon):

Forward: 5’- CTTCCAGCTCCGTGCTAAC-3’

Reverse: 5’- CGTCGCGTATTCGTTGTACT-3’

## Acknowledgments

The following kindly provided reagents used in this study: The Bloomington Drosophila Stock Center, The Vienna Drosophila Resource Center, Developmental Studies Hybridoma Bank, and Andreas Bergmann (U. Mass). Michael Sepanski (Carnegie Institution), Dongwon Lee (Duke), and Erez Cohen (University of Michigan) provided valuable technical assistance. We thank Fox laboratory members for valuable comments on the manuscript.

## Data availability

RNA-seq data will be available in NCBI Gene Expression Omnibus.

