## Supplemental Figures and Legends for "Loofah, a newly characterized adhesion protein, suppresses cell death in long-lived *Drosophila* hindgut enterocytes"

**Supplemental Data**

**5 Supplemental Figures**

**3 Supplemental Tables**

**Supplemental Table 1- RNAseq changes between young and mature adult hindgut ileums.**

**Supplemental Table 2- Primary screen data for enhancers of *hid* animal lethality.**

**Supplemental Table 3- Fly stocks used in this study.**

**Supplemental Figure Legends**

**SFig1. SDS does not cause pyloric cell death and aging does not reduce ileal EC number**
**A)** Adult pyloric epithelium from an animal fed a control diet. White= nuclei, scale bar= 25µm. **B)** Adult pyloric epithelium from an animal fed an SDS-containing diet. White= nuclei, scale bar= 25µm. **C)** Adult ileal cell number in 2d and 35d old animals. No significant difference, unpaired t-test.

**SFig2. Cell death pathway impact on midgut progenitors and hindgut pylorus, and caspase reporter data.**
**A-A')** Adult midgut ISCs and EBs expressing *esg Gal4>UAS-GFP* (green) **A** or *esg Gal4>UAS-GFP, UAS-hid,UAS-rpr* **A'** to induce cell death. White= nuclei, scale bar= 50µm. N= 6, 2 biological replicates. **B)** Adult pylorus expressing *UAS-diap1RNAi,* 4d, N=12. **C)** Adult ileum expressing *UAS-diap1RNAi,* 4d, N=12. **D-E)** Adult hindgut, caspase reporter. Magenta= PARP, Venus= GFP, Cyan= Nuclei, scale bar= 50µm. **D-D')** Adult pylorus (**D**) and ileum (**D'**), expressing *byn Gal4* (no caspase pathway transgene). **E-E')** Adult pylorus (**E**) and ileum (**E'**) 2d expressing *byn Gal4>UAS-hid*. **F)** Quantification of PARP+ nuclei, ANOVA, Tukey's, $p<0.0001$. **G)** Rescue experiments, scored for visible cell death (damage/no damage). Genotypes are indicated. At least 6 animals per conditions, 3-4 replicates.

**SFig3. Overexpression of *dronc* in the pylorus causes minor cell death.**

**A)** Adult pylorus. White=nuclei, scale bar= 50µm. **A)** Pylorus, no damage, N=24. **A')** Pylorus, 4d *bynGal4>UAS-dronc* damage, 0d recovery, N=9. **A")** Pylorus, 4d *bynGal4>UAS-dronc* damage, 6d recovery, N=17. **B)** Quantification of the extent of pyloric damage, 3-4 biological replicates, Fisher's exact test, p<0.0001.

**SFig4. Mature adult ileum ECs accumulate markers of senescence.**

**A)** 2d adult ileums. Magenta= TREDsRed, Green= γH2Av, White= Nuclei, scale bar= 50µm. **B-C)** 5d adult ileums, scale bar= 50µm **B)** Magenta= TREDsRed, White= Nuclei. **C)** Magenta= γH2Av, White= Nuclei. **D)** Percentage of TREDsRed (unpaired t-test, p<0.02) and γH2Av+ (unpaired t-test, p<0.001) cells in the adult ileum of animals 2d and 4d animals. Each data point equals one animal. From 1-2 biological replicates.

**SFig5. Loofah is an Ig domain containing protein and *loofah* RNA is efficiently reduced via RNAi.**

**A)** Loofah protein structure. **B)** RT-pPCR of salivary glands. NP5169 Gal4 was used to knockdown *loofah* in the salivary glands of wandering third instar larvae. *loofah* RNA levels are significantly reduced via RNAi (unpaired t-test, p<0.0001).

### SFig1

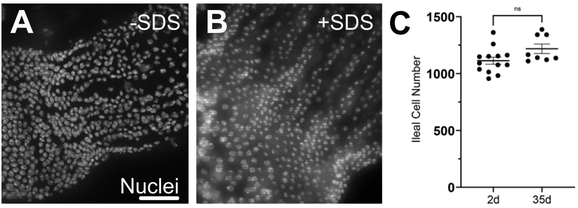

SFig2

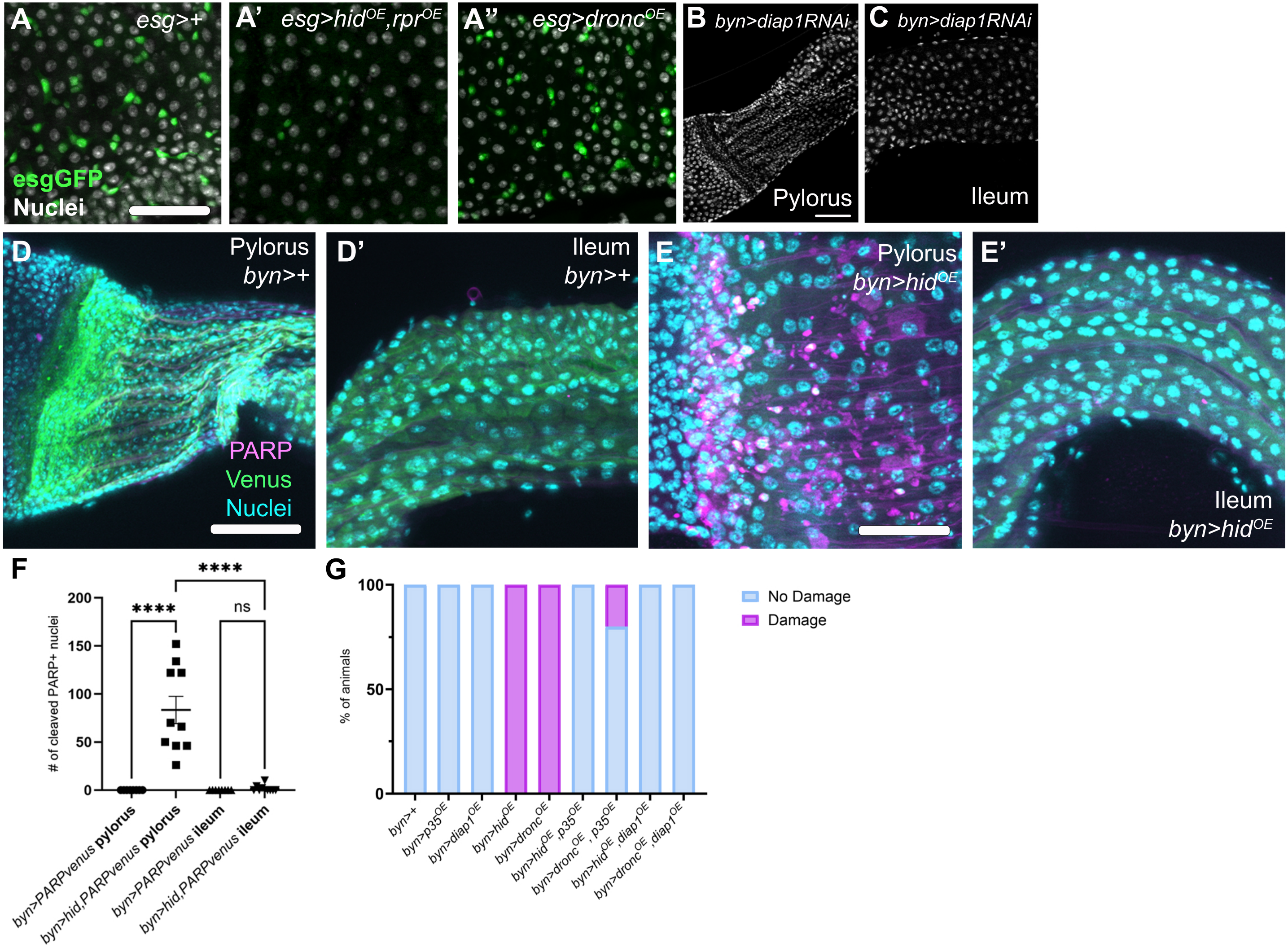
A
esg>+
esgGFP
Nuclei
A'
esg>hid$^{OE}$,rpr$^{OE}$
A''
esg>dronc$^{OE}$
B
byn>diap1RNAi
Pylorus
C
byn>diap1RNAi
Ileum
D
Pylorus
byn>+
PARP
Venus
Nuclei
D'
Ileum
byn>+
E
Pylorus
byn>hid$^{OE}$
E'
Ileum
byn>hid$^{OE}$
F
### of cleaved PARP+ nuclei
200
150
100
50
0
****
****
ns
byn>PARPvenus pylorus
byn>hid,PARPvenus pylorus
byn>PARPvenus ileum
byn>hid,PARPvenus ileum
G
% of animals
100
50
0
No Damage
Damage
byn>+
byn>p35$^{OE}$
byn>diap1$^{OE}$
byn>hid$^{OE}$
byn>dronc$^{OE}$
byn>hid$^{OE}$,p35$^{OE}$
byn>dronc$^{OE}$, p35$^{OE}$
byn>hid$^{OE}$,diap1$^{OE}$
byn>dronc$^{OE}$,diap1$^{OE}$

### SFig3

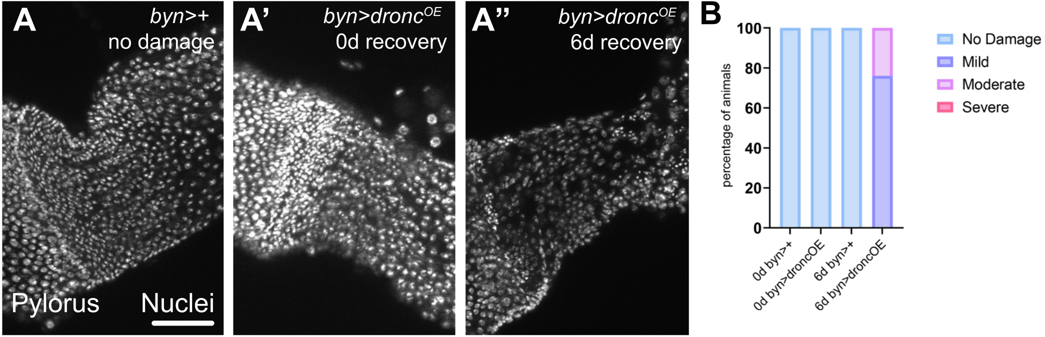
A
byn>+
no damage
Pylorus
Nuclei
A'
byn>droncOE
0d recovery
A''
byn>droncOE
6d recovery
B
100
80
60
40
20
0
percentage of animals
0d byn>+
0d byn>droncOE
6d byn>+
6d byn>droncOE
No Damage
Mild
Moderate
Severe

SFig4

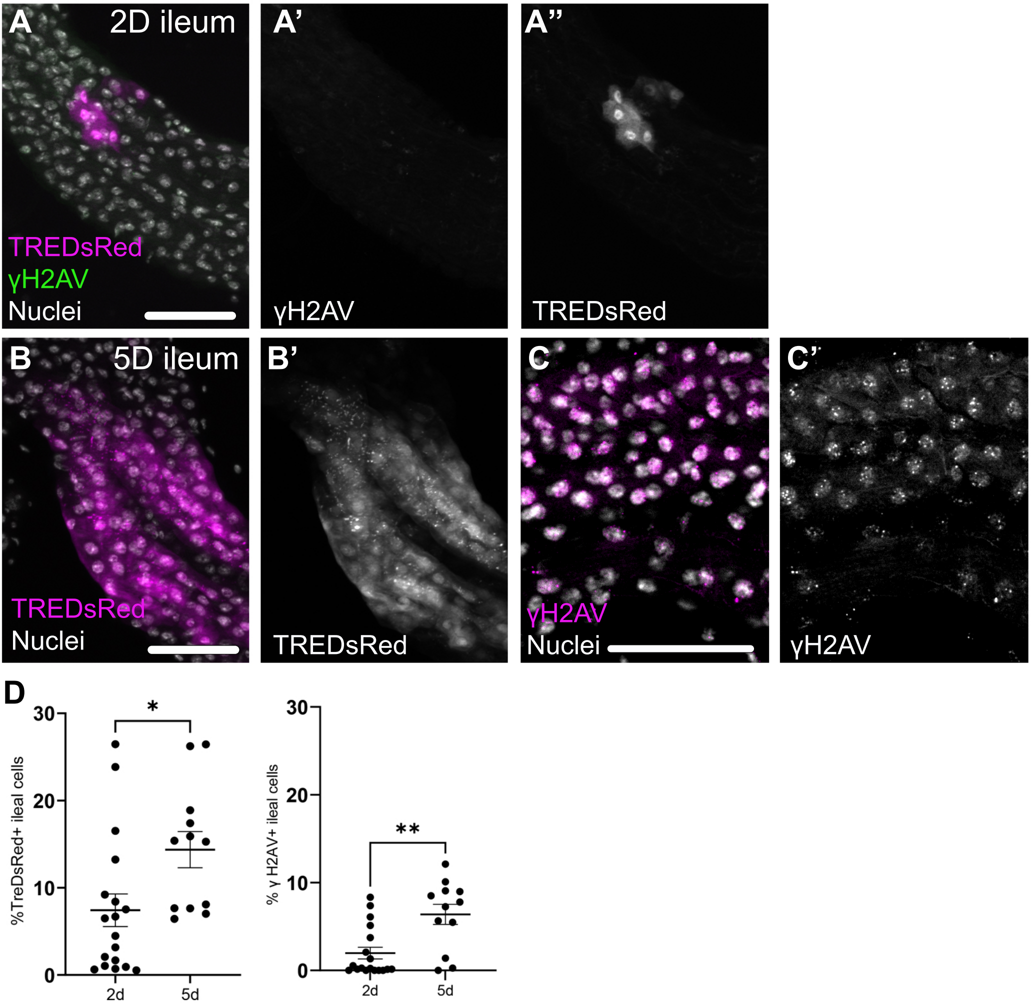
A
2D ileum
TREDsRed
γH2AV
Nuclei
A'
γH2AV
A''
TREDsRed
B
5D ileum
TREDsRed
Nuclei
B'
TREDsRed
C
γH2AV
Nuclei
C'
γH2AV
D
%TreDsRed+ ileal cells
*
2d
5d
% γ H2AV+ ileal cells
**
0
10
20
30

### SFig5

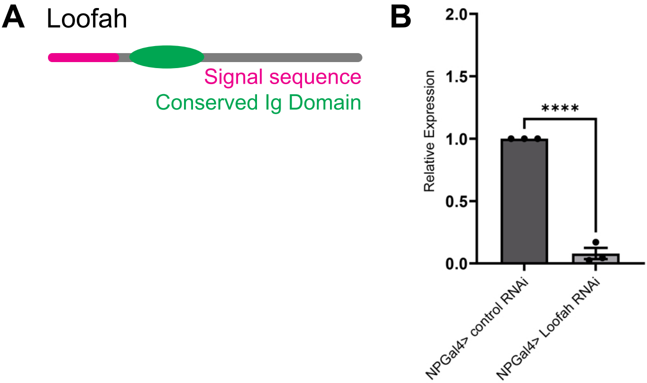
A
Loofah
Signal sequence
Conserved Ig Domain
B
Relative Expression
2.0
1.5
1.0
0.5
0.0
****
NPGal4> control RNAi
NPGal4> Loofah RNAi
